# Effects of domestication on neophobia: A comparison between the domesticated Bengalese finch (*Lonchura striata var. domestica*) and its wild ancestor, the white-rumped munia (*Lonchura striata*)

**DOI:** 10.1101/2021.03.23.436696

**Authors:** Kenta Suzuki, Maki Ikebuchi, Hiroko Kagawa, Taku Koike, Kazuo Okanoya

## Abstract

Bengalese finches (*Lonchura striata* var. *domestica*) have more complex song traits than their wild ancestors, white-rumped munias (*Lonchura striata*). Domesticated finches are likely able to allocate more resources to song development rather than allocating resources to mechanisms aimed at coping with predation, which are no longer needed under domesticated conditions. Here, we aimed to examine the effects of changes in selection pressure due to domestication on the behaviour of Bengalese finches and to contemplate the possible evolutionary mechanisms underlying these changes. To do so, we compared neophobic responses to novel-object conditions as an assessment of reactions to potential predators. We studied groups of Bengalese finches and white-rumped munias and found that Bengalese finches were more likely to eat the food provided to them under novel-object conditions. Bengalese finches had a shorter latency time to eat, and this latency time was less affected by the novel object in the case of Bengalese finches compared to white-rumped munias. Therefore, Bengalese finches have reduced neophobic responses due to domestication. The behavioural strategies of white-rumped munias appear to be more suitable for natural environments, which include unpredictable risks, whereas Bengalese finches have likely adapted their behaviour to the conditions of artificial selection.

## 1. Introduction

The Bengalese finch (BF, *Lonchura striata* var. *domestica*) is a domesticated variety of the wild white-rumped munia (WRM, *Lonchura striata*), which was imported from China to Japan approximately 250 years ago (Washio, 1996; Okanoya, 2004a; Svanberg, 2008). BFs sing phonologically and syntactically complex songs, as opposed to the stereotypical simple songs sung by WRMs (Honda and Okanoya, 1999; Okanoya, 2004a, b). BFs were domesticated for their plumage colour morphs, reproductive efficiency, parental behaviour, and ability to foster other bird species (Taka-Tsukasa, 1917), but the complex songs of BF have not been selected for in an artificial manner (Washio, 1996). Complex song as a high-quality sexual trait is believed to have evolved through the domestication process in BFs (Okanoya, 2004a, b). WRMs are thought to experience strong natural selection pressures in wild environments. Conversely, domesticated BFs experience safe and resource-rich conditions under human-controlled rearing conditions (i.e., no environmental perturbations, no predation, abundant food, and low risk of parasitism and injury). Thus, we hypothesized that the relaxation of natural selection pressures and the presence of artificial selection may have led BFs to allocate fewer resources to behaviours associated with efforts for survival and more to efforts associated with reproduction.

Domestication results in species being removed from many natural selection pressures; however, domesticated species are exposed to artificial selection pressures exerted by their captive environments and by humans (Fox, 1968; Boice, 1973; Ratner and Boice, 1975; Price, 1984). This change in selection pressure results in changes in physiological and behavioural traits (Hale, 1969; Price, 1984, Künzl and Sachser, 1999; Lepage et al., 2000). Domesticated species have reduced behavioural expressions related to ontogenetic survival, such as fear responses (Desforges and Wood-Gush, 1975; Schütz et al., 2001; Campler et al., 2009), and have enhanced behavioural traits that are not directly related to survival, such as sexual and reproductive behaviours (Künzl and Sachser, 1999). Therefore, these behavioural changes have been a major target of domestication effects.

In natural environments, birds must be vigilant of unpredictable risks such as predation when foraging or eating; this is particularly important while approaching novel conditions such as novel places, objects, and food items. Neophobia is the aversion that an animal displays while approaching novel conditions (Greenberg, 2003). In birds, the most well-established behavioural responses to novel conditions (neophobic responses) are the reactions to potential predators (Greenberg, 1990). In the wild, the presence of unpredictable risks is considered to enhance induced neophobia (Brown et al., 2020). Domesticated BFs are likely able to allocate more resources to reproduction (e.g., song development) as a trade-off for a reduction in the behaviours and strategies aimed at coping with predation (e.g., neophobic responses). In the present study, to examine the effects of changes in selection pressure due to domestication on the behaviour of BFs, and to contemplate the evolutionary mechanisms underlying these effects, we compared neophobic responses to novel conditions in BFs and their wild ancestors, WRMs.

## 2. Materials and methods

### 2.1 Birds used in this study

Seventy BFs and 58 WRMs were used in the experiment. All the birds were sexually mature. BFs (n = 10) were bought from local suppliers and others (n = 60) were bred in our laboratory. WRMs were captured in the wild in Taiwan. These captured birds were reared for more than a year in our laboratory before being used in this study. Birds were housed in a group of 8 (one group of WRM) or 10 per cage resulting in us using seven groups of BFs and six groups of WRMs for our experiments. Sex ratios of the birds that made up the groups were different for each cage (BF-1 and WRM-1: all females, BF-2, 3 and WRM-2: all males, BF-4, 5 and WRM-3, 4: 50% of each sex, BF-6, 7 and WRM-5, 6: 6 males and 4 females). Birds were housed in stainless steel cages (370 × 415 × 440 mm) within an animal-rearing room at RIKEN Brain Science Institute (BSI) and given finch seed mixture, shell grit, and vitamin-enhanced water *ad libitum*. The light cycle was kept constant at 14 h light and 10 h dark. The room was maintained at an ambient temperature of approximately 25 °C, with a humidity of 50 %. The birds were acclimatized for at least four months to the animal-rearing room before performing the experiments. All experimental procedures and the housing conditions of the birds were approved by the Animal Experiments Committee at RIKEN (#H20-2-231, #H22-2-217), and all the birds were cared for in accordance with the Institutional Guidelines of RIKEN for experiments using animals.

### 2.2 Novel-object experiment

Before the experiment, each group of birds was transferred from their home cage in the animal-rearing room to a testing cage (with the same dimensions and characteristics as the home cage) in a sound-proof box. To avoid the effects of isolation stress, experiments were conducted using the same groups of birds that were housed together during keeping. The test cage was equipped with two wooden perches and with food and water. Experiments were conducted under two conditions: the control (non-object) condition and the novel-object condition. Experiments were carried out under the different conditions on different days. All groups were first examined under the control (non-object) condition to avoid the effects of disturbance by the novel-object. To eliminate the influence of the degree of acclimation to the soundproof box and the testing cage, the number of days between the two conditions (the control and the novel-object condition) was randomly set between 1 d and 7 d in both BFs and WRMs. New food and water were provided at the beginning of the experiments. Under the novel-object condition, a novel object (a small toy dog, approximately 15 mm × 15 mm × 20 mm) was placed on the food. The behaviour of birds was recorded for 60 min using a video camera. We quantified the number of groups of birds that ate food during the observation period (at least one bird in the group) and the latency time (the time that passed before first birds started eating the food) under both conditions. If the birds did not approach or eat the food during the test period, the latency time of the birds was set at the maximum time (3600 s). All groups were used only once under both conditions. All tests were conducted between 13:00 and 15:00 to avoid other factors that could affect our results, such as diurnal changes in activity, hunger, and hormone levels.

### 2.3 Statistical analyses

The numbers of groups of BFs and WRMs that ate the food during the observation period were compared with Fisher’s exact test. The latency times to start eating the food are represented as the mean ± standard error of the mean (SEM). The latency times in BFs and WRMs were analysed using a repeated measures two-way analysis of variance (ANOVA) (birds × conditions) with post-hoc Bonferroni/Dunn tests. The difference in the effect of the novel object on the latency times (calculated as the latency times under the novel-object conditions - the latency times under control conditions) between BFs and WRMs were analysed by an unpaired *t*-test with a Welch’s correction. Values of *p* < 0.05 were considered significant. We used three different statistical software packages for analyses. Statistical analyses were performed using R statistical software (version 4.03, R Foundation for Statistical Computing, Vienna, Austria) for Fisher’s extract test, Stat View software (version 5, SAS Institute Inc., Cary, NC, USA) for repeated measures 2-way ANOVA with post-hoc Bonferroni/Dunn tests, and GraphPad Prism software (version 4, GraphPad Software Inc., San Diego, CA, USA) for *t*-tests with a Welch’s correction.

## 3. Results

### 3.1 Responses of BFs and WRMs to novel objects

We examined the effects of domestication on behavioural responses to novel objects in BFs and WRMs. Under control conditions, all groups of birds (both BFs and WRMs) ate the food at some point during the observation time (BFs: 7/7, 100 %; WRMs: 6/6, 100 %). Thus, there were no differences between BFs and WRMs (*p* = 1.00). However, under the novel-object conditions, six of the seven groups of BFs (6/7, 85.7 %) ate the food at some point during the observation time, whereas none of the WRMs groups ate the food (0/6, 0 %). Therefore, there were significant differences between BFs and WRMs (*p* < 0.01). In BFs, there were no significant differences in the number of groups that ate the food between the control and novel-object conditions (*p* > 0.90). However, WRMs were significantly less likely to eat the food under the novel-object conditions (*p* < 0.01).

### 3.2 Latency times in BFs and WRMs to start eating food under control and novel-object conditions

The latency times to start eating the food were compared between BFs and WRMs (Fig. 1). Under the control conditions, the latency times were 25.6 ± 9.0 s for BFs and 159.0 ± 46.7 s for WRMs. Under the novel-object conditions, the latency times were 1204.9 ± 524.9 s for BFs, but none of the WRMs approached the food within the observation time (3600.0 ± 0.0 s). The latency times was significantly affected by the birds (BF or WRM, *F* (1, 11) = 16.9, *p* < 0.01), conditions (novel object or control, *F* (1, 11) = 69.4, *p* < 0.0001), and interactions (birds × conditions, *F* (1, 11) = 12.7, *p* < 0.01). Latency times to eat were significantly shorter in BFs than in WRMs regardless of the presence or absence of novel objects (control conditions: *p* < 0.05, novel-object conditions: *p* < 0.01). In both species, the latency times were extended by the presence of the novel object (BFs: *p* < 0.05, WRMs: *p* < 0.0001), and the extended times were significantly longer in WRMs than those in BFs (BFs: 1379.3 ± 530.7 s, WRM: 3441.0 ± 46.7 s; *t* = 3.87, *p* < 0.01).

**Fig. 1.**
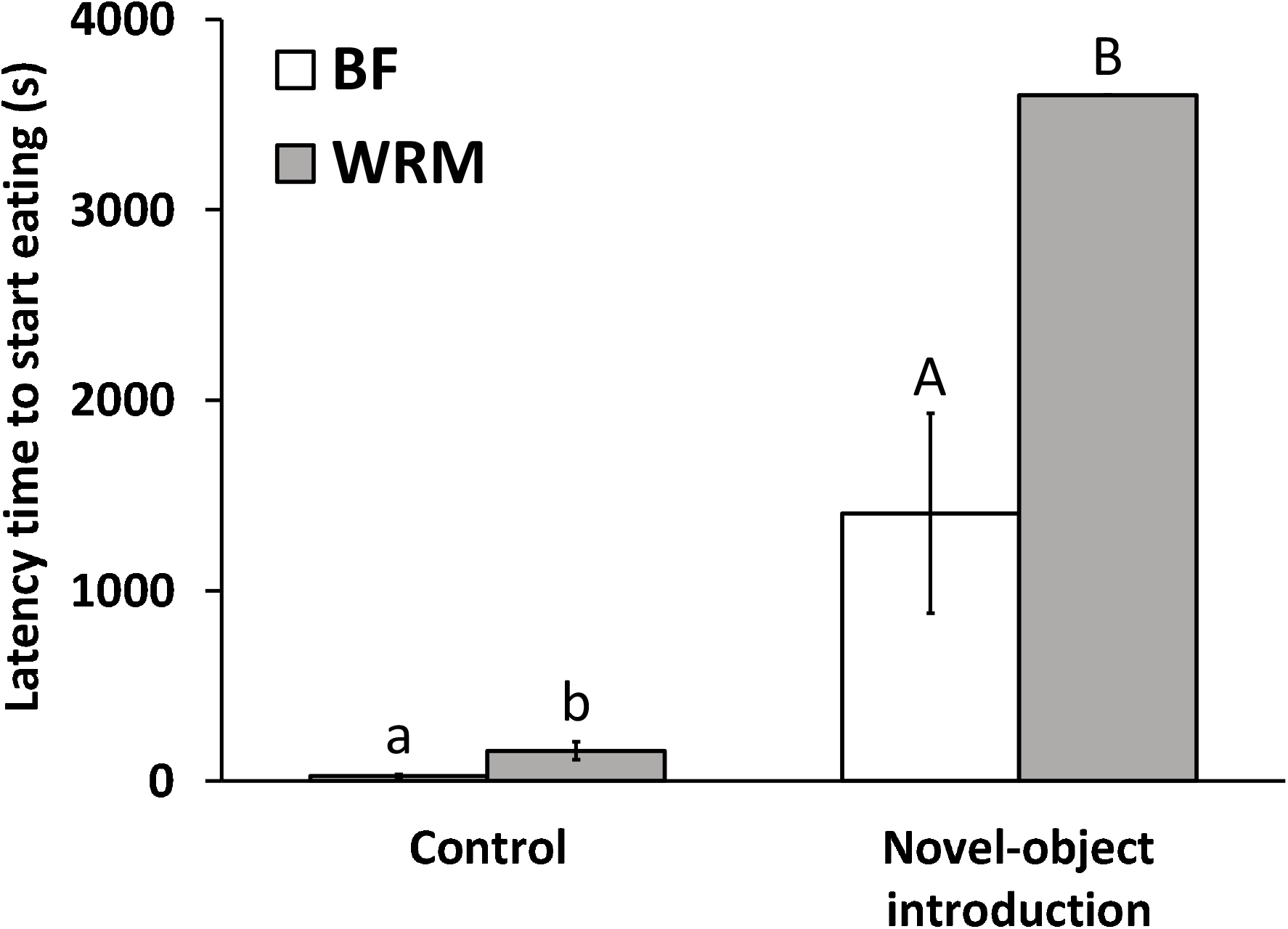
Latency times (in seconds) to start eating food in Bengalese finches (BFs, *Lonchura striata* var. *domestica*) and white-rumped munias (WRMs, *Lonchura. striata*). Bars indicate means, and vertical lines indicate standard errors of mean (SEM). BFs had significantly shorter latency times to start eating the provided food than those WRMs under both conditions (control and exposure to a novel object, i.e., a small dog toy). Latency times were extended under novel-object conditions in both BFs and WRMs. Different letters indicate significant differences between groups (a vs b: *p* < 0.05, A vs B: *p* < 0.01, a vs A: *p* < 0.05, b vs B: *p* < 0.0001).

## 4. Discussion

We compared responses to novel conditions in BFs and their wild ancestor, WRMs, and examined the effects of changes in selection pressure due to domestication. Our findings indicate that these two birds responded differently to novel-object conditions. BFs had significantly lower neophobia levels (more groups of BFs ate the food, and BFs had shorter latency times to eat under novel-object conditions) than those of their wild ancestors, the WRMs. The novel objects extended the latency times to eat in both birds, but latency times were significantly longer in WRMs than those in BFs. Therefore, it is likely that BFs have reduced neophobic responses due to domestication. Under control conditions, the proportion of the groups that ate the food was the same in BFs and WRMs. However, BFs ate the food sooner than WRMs. It is probable that the WRMs were more cautious about the experimental situation (the soundproof box is different from the WRMs’ normal environment). In a previous report, male WRMs had higher faecal corticosterone concentrations than those in male BFs (Suzuki et al., 2012; 2014a, b). Therefore, WRMs are considered to have a higher level of baseline vigilance than that in BFs. This strain difference with respect to vigilance seems to have been further increased under novel-object conditions.

Our results are similar to those of a previous study that found that domesticated ducks (*Anas platyrhynchos* var. *domesticus*) had lower levels of neophobia than those of wild mallards (*Anas platyrhynchos*) (Desforges and Wood-Gush, 1975). In addition, laboratory mice (*Mus musculus*) had lower levels of neophobia than those in wild mice (Meddock and Osborn, 1968). In a comparison of dogs (*Canis lupus familiaris*) and wolves (*Canis lupus*), dogs had lower levels of neophobia than those in wolves, but dogs had an overall lower interest in novel objects (Moretti et al., 2015). In the present study, BFs showed a fear response to novel objects and were considered to have no diminished interest in novel objects.

This study was conducted using groups of birds, and, as such, there may have been a social effect. Because it was difficult to identify individuals in this study, the analysis was conducted as a group-by-group comparison, and it was therefore not possible to take into account the differences in the responses of each individual. Wolves and dogs spend more time approaching a novel object in groups than as individuals, so risk sharing may increase vigilance (Moretti et al., 2015). In contrast, in the zebra finches (*Taeniopygia guttata*), existence of conspecifics can modulate (decrease) the responses to stressors (e.g., novel stimuli), known as “social buffering” (Coleman and Mellgren, 1994; Emmerson and Spencer, 2017). Responses to novel objects may also be influenced by the identities of the group members. In the Gouldian finches (*Chloebia gouldiae*), shy birds took more risks when they were paired with bold partners, and bold birds took fewer risks when they were paired with shy partners (King et al., 2015). Since this experiment was conducted using groups, there may have been an effect of risk sharing or social buffering. Therefore, in experiments conducted using groups, individual differences may be reduced, and more of the characteristics of the groups may be expressed. Conversely, in experiments conducted using individuals, there is a high possibility that individual differences will be noticeable. Future studies on BFs and WRMs should take into consideration the reactions of each individual bird to novel-object conditions. In a previous study, we compared the tonic immobility response, which can be used as a measure of fear responses, between individual BFs and WRMs and found that BFs had lower fear responses (Suzuki et al., 2013). Therefore, we believe that the fear response is lower in BFs than that in WRMs, even on an individual basis.

In this study, the effects of sex ratio and environmental or rearing conditions (bought from local supplier, wild-caught, or bred in the laboratory) were ruled out because the groups in the cages were neither separated by sex nor by environmental or rearing conditions. Moreover, previous studies have shown that sex and environmental or rearing conditions of the white-rumped munia (captive-born, bought from a supplier, or captured) did not affect the fear response (tonic immobility reactions) (Suzuki et al., 2013). Additionally, the conditions under which Bengalese finches and white-rumped munias were bred or reared did not affect the corticosterone levels, which is known to affect fear responses (Suzuki et al., 2012). All the munias used in this experiment were wild-caught; however, differences in rearing conditions have not been an issue in previous experiments. Therefore, the magnitude of the impact of these factors is small, but it must be considered in future research.

We hypothesized that BFs may allocate more resources to reproduction (e.g., song development) as a trade-off for a reduction in the behavioural strategies aimed at coping with predation (e.g., neophobic responses). The behavioural strategies of WRMs appear to be more suitable for natural, wild environments, which include unpredictable risks (e.g., predation risk), whereas BFs have likely adapted their behaviour to the conditions of artificial selection. Singing is important for reproduction but can increase susceptibility to predators. Birds are able to respond to increased predation risk by reducing singing, and it has been found that in areas with increased perceived predation risk, fewer songs are produced per bird (Abbey-Lee et al., 2016). Continuous auditory feedback is necessary to maintain the complex songs of the BF (Okanoya and Yamaguchi, 1997), and domesticated BFs might be able to allocate more time to singing and song maintenance due to the reduced need for neophobic responses. Songs incur a cost in their development and maintenance linked to corticosterone levels (Gil and Gahr, 2002; Okanoya, 2004a, b; Wada et al., 2008; MacDougall-Shackleton et al., 2009). Administration of corticosterone during the developmental period enhances the fear response (including the neophobic response) but reduces song complexity (Spencer et al., 2003; Buchanan et al., 2004; Emmerson and Spencer, 2017). Chronic stress and corticosterone treatment enhance dendritic expansion in neurons of the basolateral amygdala and increase anxiety as a function of that brain region (Vyas et al., 2002; Vyas and Chattarji, 2004; Mitra et al., 2005; Mitra and Sapolsky, 2008). In contrast, chronic stress and corticosterone administration in juvenile and adult birds reduce the volume of song nuclei and the numbers of new and matured neurons in the HVC (Buchanan et al., 2004; Newman et al., 2010). Both the amygdala and the song nuclei have two receptors (mineralocorticoid and glucocorticoid receptors) for corticosterone, and different levels of their expression might cause different site-specific effects (Matsunaga et al., 2011; Suzuki et al., 2011; Suzuki et al., 2014a). Therefore, low levels of corticosterone in BFs (Suzuki et al., 2012; Suzuki et al., 2014b) are considered to be required for the development of low neophobic responses and complex songs.

In summary, our results suggest that the domestication process led to differences in the responses of BFs and WRMs to novel conditions. The behavioural strategies of WRMs seem to be suitable for the natural environment, which includes unpredictable risks, whereas BFs might have adapted their behaviours to the conditions of captivity and artificial selection. BFs are likely able to allocate the resources that would be needed in the wild to cope with predators to song development, due to domestication. In addition, there were variations in latency times to eat under novel object conditions in BFs, which indicate individual differences in neophobic responses. Although this study was a group-by-group examination, future studies would need to focus on individual differences and examine the trade-off between personality traits (such as neophobia) and reproductive traits (such as song complexity). This is the first study to evaluate neophobia in Bengalese finches and white-rumped munias, and also explore the evolutionary mechanisms of behavioural changes in domesticated animals, including the evolution of complex songs.

## Abbreviations

BF: Bengalese finch
WRM: white-rumped munia.

## Acknowledgements

We thank members of the Biolinguistics team (RIKEN BSI, Saitama, Japan) for animal care. This study was supported by RIKEN BSI and Japan science and technology agency (JST) - the exploratory research for advanced technology (ERATO) research funding.

## Author contributions

**Kenta Suzuki:** Conceptualization, Data curation, Formal analysis, Investigation, Methodology, Visualization, Writing - original draft; **Maki Ikebuchi:** Conceptualization; **Hiroko Kagawa:** Conceptualization, Writing - review & editing; **Taku Koike:** Data curation, Investigation; **Kazuo Okanoya:** Conceptualization, Funding acquisition, Supervision, Writing - review & editing.

